# Exploring changes in social spider DNA methylation profiles when succumbing to infection in CpG, CHG, and CHH contexts

**DOI:** 10.1101/2024.05.21.595160

**Authors:** David N. Fisher, Jesper Bechsgaard, Trine Bilde

## Abstract

Living at high density and with low genetic diversity are factors that should both increase the susceptibility of organisms to disease. Therefore, group living organisms, especially those that are inbred, should be especially vulnerable to infection and therefore have particular strategies to cope with infection. Phenotypic plasticity, underpinned by epigenetic changes, could allow group living organisms to rapidly respond to infection challenges. To explore the potential role of epigenetic modifications in the immune response to a group-living species with low genetic diversity, we compared the genome-wide DNA methylation profiles of five colonies of social spiders (*Stegodyphus dumicola*) in their natural habitat in Namibia at the point just before they succumbed to infection to a point at least six months previously where they were presumably healthier. We found increases in genome- and chromosome-wide methylation levels in the CpG, CHG, and CHH contexts, although the genome-wide changes were not clearly different from zero. These changes were most prominent in the CHG context, especially at a narrow region of chromosome 13, hinting at an as-of-yet unsuspected role of this DNA methylation context in phenotypic plasticity. However, there were few clear patterns of differential methylation at the base level, and genes with a known immune function in spiders had mean methylation changes close to zero. Our results suggest that DNA methylation may change with infection at large genomic scales, but that this type of epigenetic change is not necessarily integral to the immune response of social spiders.

## Introduction

Pathogens and parasites are a key source of mortality and fitness reduction in essentially all organisms. Consequently, pathogens and parasites are expected to be key drivers of evolution, adaptation, and speciation (Van Valen, 1973). Parasites and pathogens are expected to play especially key roles in group living species, as disease can spread more easily when organisms live at high densities and frequently interact (Cremer *et al*., 2007; Nunn *et al*., 2015). As such, group living organisms are expected to show numerous adaptations to parasites and pathogens (Schmid-Hempel, 1988). Variation among individuals in their genes is the raw material upon which selection acts to generate adaptive change and may help protect populations from disease by increasing the chances at least some individuals will be partially resistant (Altermatt and Ebert, 2008). However, genetic variation in fitness-relevant traits is expected to be low, and in some group-living organisms individuals may be related due to inbreeding, hence lowering genetic diversity within the group even further (Hatchwell, 2010; Settepani *et al*., 2017). Social groups therefore present a challenge: how do they cope with exposure to parasites and pathogens when living at high densities and when genetic variation among group members is low?

Key candidate mechanisms through which populations with low genetic diversity can still show rapid responses to environmental threats are epigenetic modifications such as histone modification and DNA methylation (Flores *et al*., 2013). Epigenetic mechanisms can act within an organism’s life to alter gene expression and therefore phenotypes without altering the genetic code (Ledon-Rettig *et al*., 2012), allowing much more rapid responses to immune challenges compared to adaptive change though changes in allele frequencies across 10s of generations. For example, in both silkworm (*Bombyx mori*) and a mosquito (*Anopheles albimanus*) the immune response is modulated by DNA methylation in the midgut (Wu *et al*., 2017; Claudio-Piedras *et al*., 2020). DNA methylation occurs when the carbon-5 position of a cytosine base gains a methyl group. DNA methylation is one of the more commonly studied epigenetic mechanisms, and in mammals and plants is known to reduce expression of genes when bases within genes promoters are methylated (Bird, 2002; He *et al*., 2011; Elhamamsy, 2016). Further, in both plants (Deleris *et al*., 2016) and mammals (Mostoslavsky and Bergman, 1997; Morales-Nebreda *et al*., 2019) DNA methylation plays a key role in the immune response to both bacterial and viral pathogens. The role of DNA methylation is less clear in arthropods and other invertebrates however, with DNA methylation being associated with both increased, decreased, and stabilised gene expression (Richard *et al*., 2021; Duncan *et al*., 2022), and so whether DNA methylation plays an important role in the immune response requires investigating.

Social spiders are an excellent model system to study how group living species resist parasites and pathogens as they live in dense aggregations with constant social interactions among individuals, creating the perfect conditions for disease transmission. Further, social spiders are highly inbred, meaning groups have incredibly low genetic diversity (Settepani *et al*., 2017), and thus in theory should be especially susceptible to pathogens. This is supported by the recent findings that social spiders have impaired immune responses compared to their non-social sister-species (Bechsgaard *et al*., 2022), and that nests do not persist very long likely due to diseases (Busck *et al*., 2022). Despite this, populations of social spiders persist. Comparative genomic approaches have recently shown that genes rapidly evolving in social spiders compared to their solitary sister species include those important in immune function (Tong *et al*., 2020), indicating this is the ideal taxon to explore how parasites and pathogens drive evolution in group-living organisms. Further, the genome of the social spider *Stegodyphus dumicola* has relatively high methylation level, and the patterns are consistent with a role of DNA methylation in local adaptation (Liu *et al*., 2019; Aagaard *et al*., 2022). Genes that are methylated have higher and more stable expression patterns among individuals (Liu *et al*., 2019), suggesting DNA methylation could be a mechanism underpinning changes in gene expression and therefore phenotypic plasticity. However, the functional role of DNA methylation in this taxon is still opaque. The spider immune response involves phagocytosis and encapsulation, production of effector molecules such as hemocyanin and phenoloxidase, and a clotting cascade (as well as constitutive responses; Kuhn-Nentwig and Nentwig, 2013; Bechsgaard *et al*., 2016)( all of which could be regulated by gene expression and therefore DNA methylation.

Here we aim to characterise how patterns of DNA methylation change in *S. dumicola* shortly before nest extinction, which is associated with a high bacterial load (Busck *et al*., 2022). Given the high bacterial load typically present in nests before they die off, we can expect them to be mounting an immune response compared to when they are healthy. By comparing DNA methylation profiles of a nest when it is healthy and when it is about to die off, we can therefore determine the role of DNA methylation in the immune response. We performed this comparison at a range of scales (base, gene, chromosome, whole genome) to gain a holistic understanding of how DNA methylation changes in response to infection.

## Methods

### Sample collection

*Stegodyphus dumicola* is a social spider found in central and southern Africa (Majer *et al*., 2013). Individuals live in groups of tens to hundreds of individuals, typically very highly related to one another (Lubin and Bilde, 2007; Settepani *et al*., 2017). They cooperate during reproduction and foraging, including transferring digestive fluids during communal external digestion of prey (typically flying insects) leading to highly similar microbiomes among nest mates (Busck *et al*., 2020; Rose *et al*., 2023). Such close associations also create the ideal conditions for the transmission of pathogens, and indeed it is thought both bacterial and fungal infections are a key source of mortality (McEwen *et al*., 2020; Busck *et al*., 2022). As part of a larger study, we located five nests on farmland in northern Namibia (Busck *et al*., 2022). After locating the first of these nests in April 2017, we returned approximately every three months to locate new nests, record the survivorship of existing nests, and sample three individuals from each nest. Individuals were extracted from each nest by vibrating the capture web or gently squeezing the nest until spiders emerged, and then were placed individually in microcentrifuge tubes with ATL buffer (Qiagen, Hilden, Germany) and transferred to a portable freezer. Here we use the three spiders from the last time point a nest was sampled before it was no longer observed or was observed to be devoid of live spiders (hereafter “dying”) and compare them to three spiders from a time point 6-12 months prior to that (hereafter “alive”; see Fig. 1). We chose the alive time point based on whether the spiders were a similar size to the three dying spiders, as this should indicate they are a similar age, although we were limited by sample availability as to how well we could match sizes (see Table S1 for sample details, including estimates of bacterial load for some samples). Given nest extinctions are associated with a large increase in bacterial load both in the last observation before death and three months before that (Busck *et al*., 2022), by comparing dying spiders with those 6-12 months prior of the same nest we aim to conduct a paired analysis of infected compared to not-infected nests. A paired design is preferable to comparing a group of infected nests and a group of healthy nests as among-colony variation in methylation profiles is large in eusocial hymenoptera (Libbrecht *et al*., 2016; Marshall *et al*., 2019; Cardoso-Júnior *et al*., 2021) and a similar patterns in social spiders would limit our ability to detect a difference between the infected and non-infected groups.

**Figure 1.**
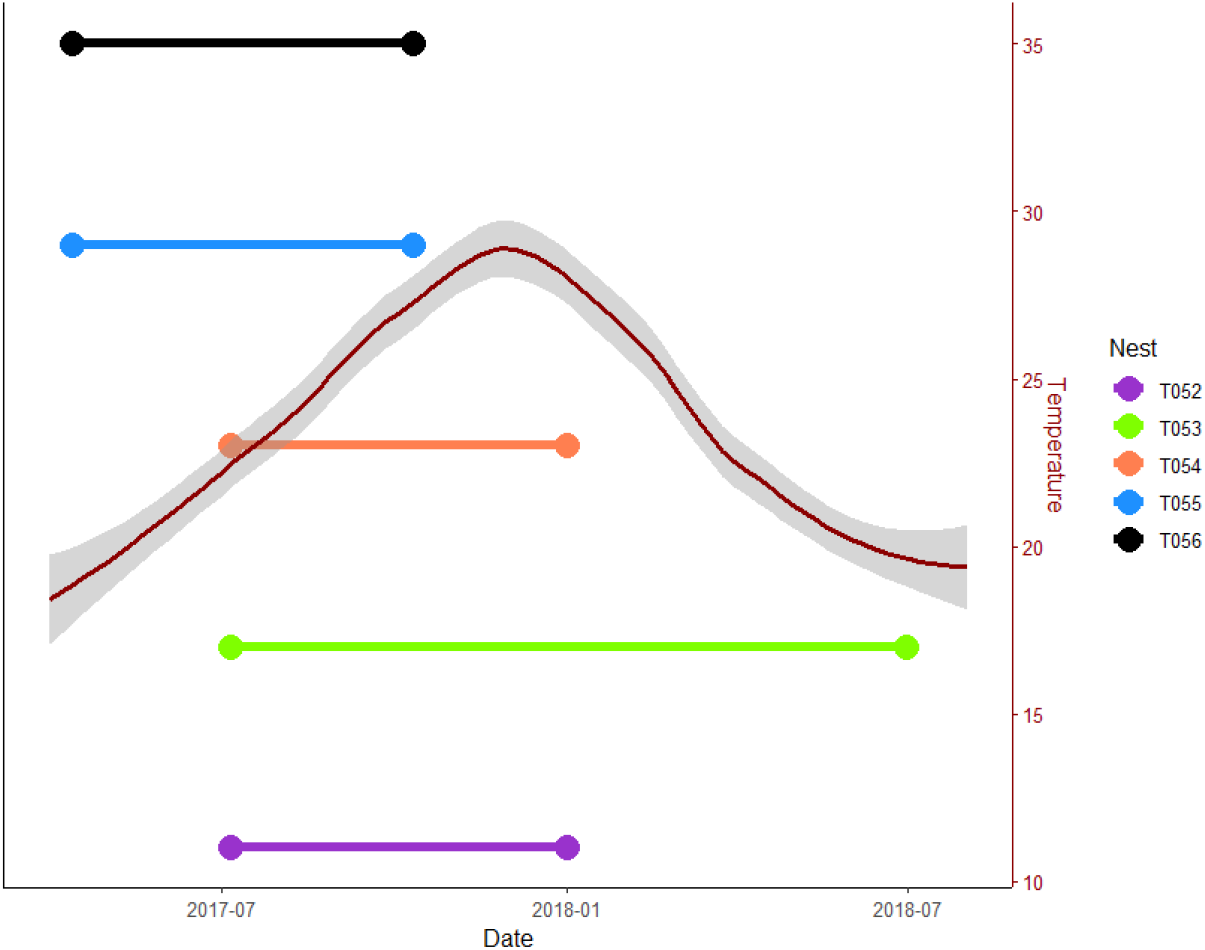
Time points our samples took place, with a smoothed line in red showing the mean temperature (and 95% confidence intervals in grey) during that time to give an idea of season. The earlier time point for each nest is the alive time point, while the latter is the dying time point. Positions of the nests on the y-axis are just for illustration.

### DNA extraction and sequencing

We extracted DNA using a DNeasy Blood and Tissue kit from Qiagen (Hilden, Germany). We pooled the DNA from the three spiders from each nest at each time point and subjected each of these ten samples to bisulfite treatment. Bisulfite treatment converts unmethylated cytosines to uracils, while methylated cytosines are unconverted (Cokus *et al*., 2008). Subsequent sequencing and mapping to a reference genome (Ma *et al*., 2024) then allows one to determine the location of each methylated cytosine. We used DNBSEQ-400 for 100 bp paired-end sequencing (BGI, Hong Kong). To measure the bisulfite conversion error rate we used λDNA, which indicated that >99% of unmethylated cytosines were successfully converted. After sequencing adaptor sequences, contamination, and low-quality reads were removed from the raw data.

### DNA methylation quantification

We identified methylated and unmethylated cytosines using Bismark v 0.23.1 (Krueger and Andrews, 2011). First, we indexed the reference genome via bowtie2 with “bismark_genome_preparation”, using the default parameters. Following this, we used the function “bismark” to map the bisulfite converted sequencing reads for a given sample, and the alignment was used to call the methylation status of all cytosines across the genome (default parameters). A small number of alignments can be duplicated, so we removed these using “deduplicate_bismark”. Following this, we extracted the coverage and methylation status for each cytosine base covered by at least one read using “bismark_methylation_extractor”, indicating paired ends (-p), including only a single read from overlapping sections (--no_overlap), and including cytosines in all contexts (--comprehensive). We did this as, while CpG DNA methylation is the most common, both generally and in the *S. dumicola* genome (Liu *et al*., 2019), important patterns can appear in CHG and CHH contexts; (Dubin *et al*., 2015; Hämälä *et al*., 2022). We then created M-bias plots to determine if there were biases in the methylation calls in our reads (Hansen *et al*., 2012). We observed a bias towards low methylation percentages in the first two bases of the 3’ end, which is often found and results from the sequence of the adapter used. We also observed high variability in the first five bases of the 5’ prime end (Figs. S1-10). As such, we repeated the bismark_methylation_extractor call with --ignore 5 and --ignore_r2 2 to remove these areas of bias. We additionally generated a “bedgraph” format output file containing the position of each cytosine, the coverage, the number of methylated cytosines, the number of unmethylated cytosines, and the percentage of reads that were methylated, for cytosines in each of the CpG, CHG, and CHH contexts. We excluded cytosines from this output file if they had a coverage of less than five reads, as their methylation status would be more uncertain, and those with more than 29 reads, which likely indicated some sort of PCR bias (99.9% of bases had fewer than 30 reads). These steps resulted in thirty output files, three for each of the ten samples.

### DNA methylation analysis

We first determined if the overall percentage of methylated cytosines across the entire genome changed between the alive and dying time points. To do this we followed Liu et al. (2019) in defining each cytosine in our output files as either methylated or not using a binomial test. For each cytosine, the number of methylated reads was entered as the number successes and the coverage as the number of trials, and we tested whether the observed proportion methylated was greater than 0.01 (the error rate estimated with the λDNA, see above). We corrected all p-values using the Benjamini-Hochberg procedure for multiple tests, and defined bases with a p-value lower than 0.05 as methylated, and all others as not. For each sample we calculated the genome-wide mean methylation proportion and tested whether there was a clear difference between the alive and dying time points using a Wilcoxon signed rank test. We predicted that overall methylation would increase if methylation was integral to fighting the infection as there would be increased methylation to increase gene expression of relevant pathways. We also calculated the mean methylation proportion per chromosome, and the mean methylation proportion per 1000 bp region, and used Wilcox-tests to determine if counts in these regions were different between the alive and dying time points and F tests to determine if the variances among these regions were different between the alive and dying time points. We predicted that both the mean and the variance would be greater in the dying group than the alive group, as differentially methylating immune genes would lead to overall-higher but also a more heterogeneous patterns of methylation. Next, we examined the methylation response at the base and gene level using the R package “methylKit” v 1.20.0 (Akalin *et al*., 2012). For this we returned to the original bedgraph output files. First, we filtered out bases with fewer than 5 or more than 30 reads, but otherwise we retained information on number of reads and percentage that were methylated, rather than assigning a cytosine the status of methylated or not. We also only used bases that were read in at least three samples i.e., their methylation state was observed for at least two different nests.

Base-level analyses: We performed an F-test with the McCullagh and Nelder (1989) correction for overdispersion to test for bases that were consistently differentially methylated (either hypo or hypermethylated) between the alive and dying time points, treating each nest as a biological replicate rather than pooling them, including nest identity as a categorical effect, and calculated the mean methylation difference using read coverage as weights. We identified bases that were clearly differentially methylated (DMBs) as those with a change in methylation percentage of at least five percentage points, and with a p-value, after a Benjamini-Hochberg correction for false discovery rate, of ≤0.05. In the CpG context this corrected p-value corresponds to an uncorrected p-value of 1.3086 x 10^-7^, only marginally higher than the commonly-used cut off of 5 x 10^-8^ (recently reviewed by Chen et al., 2021; note that the *S. dumicola* genome is of similar size to the human genome; Risch and Merikangas, 1996). We excluded bases on scaffolds 6 and 16 as these are not genuine chromosomes. We then associated the DMBs with an annotated genome using the R packages “plyranges” (Lee *et al*., 2019) and “Granges” (Lawrence *et al*., 2013). This allowed us to determine the genome features DMBs tended to be in e.g., regions of coding or repetitive DNA.

Gene-level analyses: To test which genes were differentially methylated, we used the original bedgraph output files and summarised the number of methylated and unmethylated cytosines by gene (as defined by the genome annotation file) using the “regionCounts” function in methylKit. We then filtered to remove all genes with more than 100,000 cytosines as these were unusually high (between 95 and 99^th^ percentile and above) and might represent a PCR bias. We then removed all genes that did not feature in at least three samples, as we did for the base-level analysis. We then performed an F-test with the McCullagh and Nelder correction for overdispersion to test for genes that were differentially methylated across samples. This analysis takes into account the total methylated and unmethylated cytosines in each gene (similar to a weighted methylation level; Schultz *et al*., 2012) when comparing the alive and dying time points. We corrected each p-value for false discovery rate and reported all those genes with a corrected p value of less than 0.05. We performed this for all colonies in the CpG context, but due to the lower rates of methylation we could not do this for the CHH context. In the CHG context, we had to exclude the “alive” sample for nest T54 and the “dying” sample for nest T55 due to no overlap between read cytosines and genes in the annotation file, and so the analysis took place on eight samples, four in each condition.

Finally, to determine if genes with a suggested immune function were differentially methylated between the alive and dying time points, we compiled a list of candidates based on immune genes identified as being present in the congeneric social spider *S. mimosarum* by Bechsgaard *et al*. (2016) and in the fellow Araneomorphid *Parasteatoda tepidariorum* by both Bechsgaard *et al*. and Palmer and Jiggins (2015; their Table S5). The FlyBase IDs for these genes (46 in total) are listed in Table S2. We used the program BLAST and the command “tblastx” to identify which of these genes were present in our *S. dumicola* genome. We then used the methylation difference % per gene calculated for the gene-level analysis for these immune genes and tested whether these gene-level scores were different from zero using a two-tailed t-test.

## Results

We obtained on average 249,287,506 alignments per sample, of which on average 2.244% (5,591,062 per alignment) were duplicates and so removed. On average across all ten samples (five nests, both alive and dying) each sample contained 6,851,906,773 cytosines; 1,124,453,155 in the CpG context, 1,244,000,324 in the CHG context, and 4,145,816,481 in the CHH context (some cytosines could not be assigned a context). The mean percentage of methylated cytosines in each was 10.22%, 0.72%, and 0.99% for the CpG, CHG, and CHH contexts respectively (114,911,915, 8,990,689, and 34,680,101 methylated cytosines in the CpG, CHG, and CHH contexts respectively).

### CpG

Three colonies showed an increase in genome wide methylation rate, while two showed essentially no change (Fig. 2a) meaning there was no clear change overall (paired Wilcoxon signed-rank test, V = 3, p = 0.313). Chromosomes showed consistent rates of methylation, with increases across all nests when the nest was dying compared to when alive (Fig. S11; paired Wilcoxon signed-rank test, V = 240, p < 0.001). There was no difference in the variance of methylation rates at the chromosome level between the alive and dying samples (variance alive = 0.0001, variance dying = 0.0001, F-test, F = 0.996, df = 69, p = 0.988). There was a small increase in mean methylation rates from alive to dying at the level of 1000 base pairs (Wilcoxon signed-rank test, V = 1.058 x 10^14^, p = 0.001) and a small decrease in the variance in methylation rates (variance alive = 0.1521, variance dying = 0.1517, F-test, F = 1.002, df = 14,545,555, p < 0.001) but this is a very small difference made clear by the very large sample size. Note the much greater variance in methylation rates at the 1000 base pair level is because many sets of 1000 base pairs have no methylation at all.

**Figure 2.**
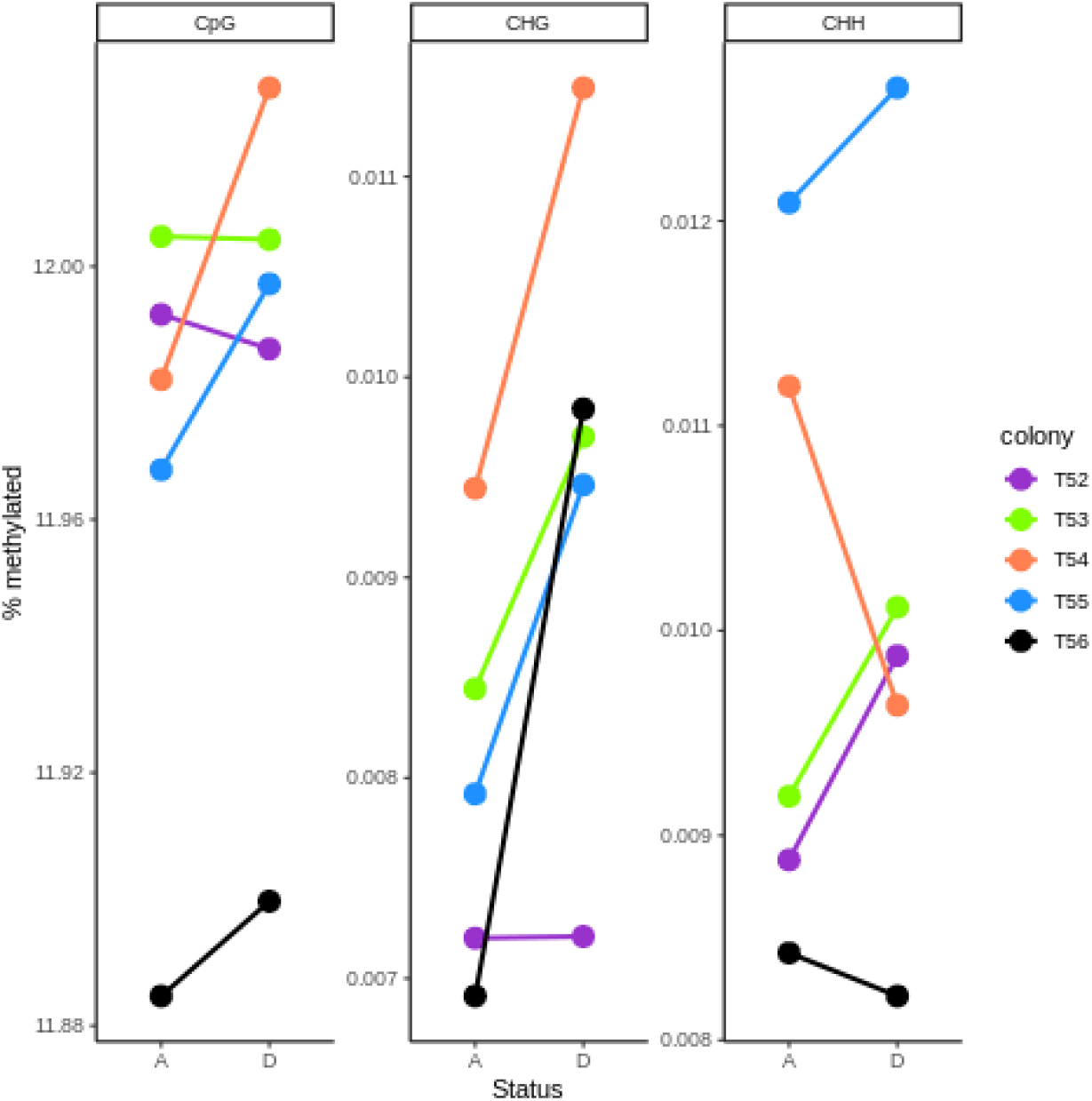
Change in genome-wide methylation percentage of cytosines in each context for nests between being alive (A) and when dying (D).

We identified 19 DMBs (differentially methylated bases) across all nests in the CpG context, eight hypermethylated and 11 hypomethylated (Figs. 3a & 3b). Of the hypermethylated DMBs, seven were in regions of repetitive DNA, with two LTR retrotransposons (Gypsy, BEL), three LINE retrotransposons (all I-Jockey), one DNA transposons TcMar) and two unclassified repeats. The remaining hypermethylated DMB was in a gene intron. Of the hypomethylated DMBs, six were in unclassified repeats in regions of repetitive DNA. The remaining hypomethylated DMBs were in gene introns.

**Figure 3.**
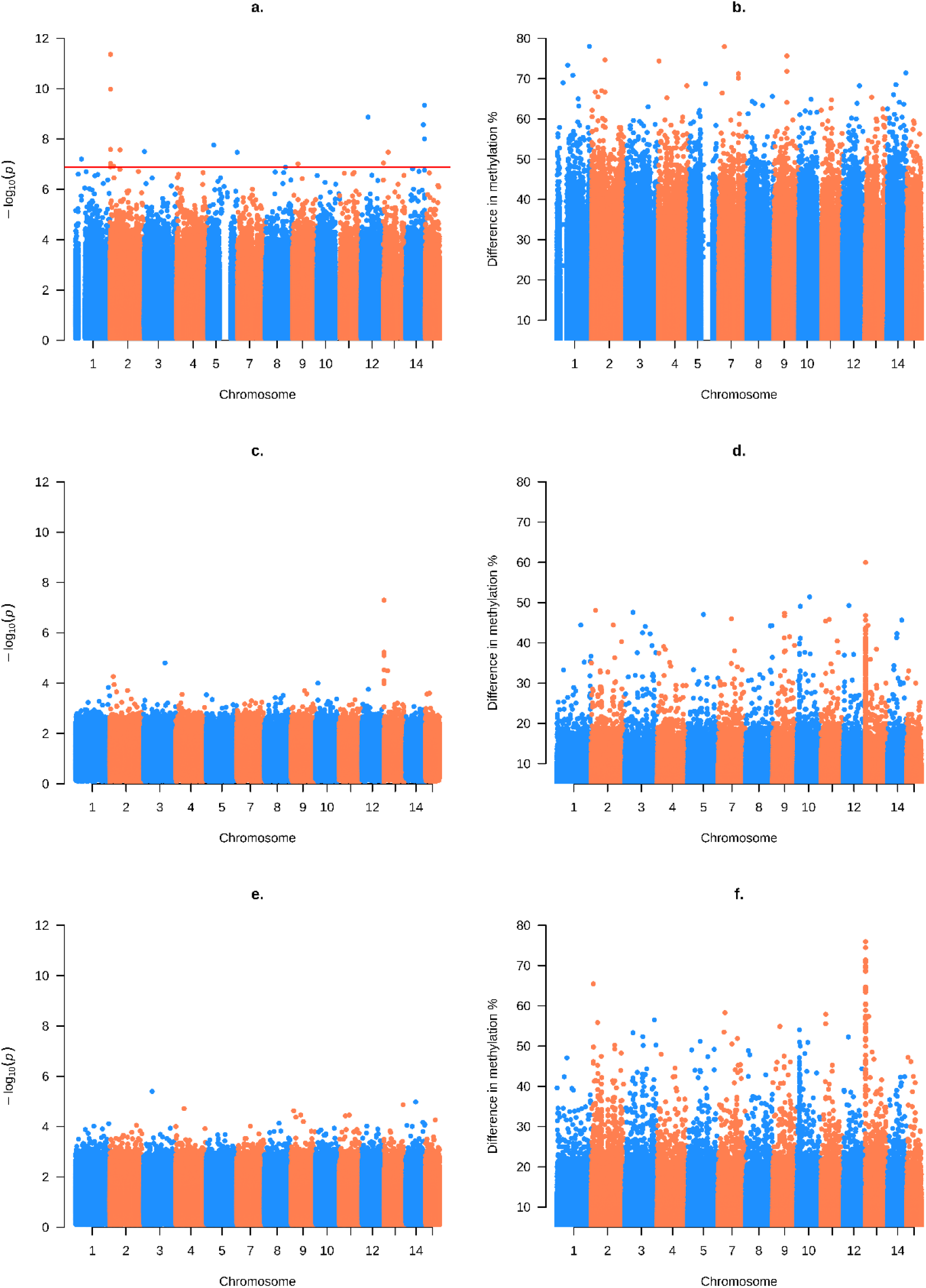
Manhattan plots of differentially methylated bases in the CpG context (a-b), the CHH context (c-d) and the CHH context (e-f). The left column (a, c, e) show -log_10_ of p values for the statistical test, while the right column (b, d, f) shows the percentage difference in DNA methylation. The red line in a. indicates where the p value, after a correction for the false discovery rate, was < 0.05 and is our threshold for a clear difference in methylation. Note no bases passed the threshold for statistical significance in the CHG and CHH contexts The use of blue and orange highlights each chromosome.

Our analysis of weighted methylation levels per gene revealed a single gene in the that was differentially methylated between alive and dying the in CpG context. This gene had a weighted methylation increase of 3.690 (q-value = 0.016, p (fdr) = 0.017), and so was hypermethylated in the dying colonies. Using the NCBI “blastp” tool at https://blast.ncbi.nlm.nih.gov/Blast.cgi we searched the gene’s sequence and found it was 99.59% identical to a heparan-sulfate 6-O-sulfotransferase 2 detected from our study species’ congener *S. mimosarum* (accession number: KFM63315; no other matches were above 88% identical).

The sequences of the 46 putative immune genes we selected were detected 1465 times in the genome. We had methylation changes in at least three samples in the CpG context of 7,417 of these, with a mean methylation difference of 0.259% (min = -8.284, max = 7.925). The mean is very close to zero, but significantly different due to the large sample size (t_7560_ = 22.709, p < 0.001, 95% CIs = 0.237 to 0.282).

### CHG

All colonies showed an increase in genome wide methylation rate when dying (Fig. 2b) although this was not clearly different from zero, presumably due to the small sample size (Wilcoxon signed-rank test, V = 0, p = 0.063). All chromosomes showed increases from alive to dying (Fig. S12; paired Wilcoxon signed rank test, V = 444, p < 0.001). The variance of methylation rates at the chromosome level was higher in the dying compared to the alive samples (variance alive = 1.511 x10^-10^, variance dying = 3.618 x10^-10^, F-test, F = 0.417, df = 69, p < 0.001). Both the mean and the variance at the level of 1000 base pairs was also higher in the dying compared to alive colonies (Wilcoxon signed rank test, W = 1.066 x 10^14^, p < 0.001; variance alive = 7.342x10^-6^, variance dying = 9.750x10^-5^, F-test, F = 0.753, df = 14,592,315, p < 0.001).

No cytosines were differentially methylated after correcting for multiple testing in the CHG context (Fig. 3c). There was however a clear spike in the p values (Fig. 3c) and % differences in methylation (Fig. 3d) at chromosome 13, approximately between bases 704,000 and 821,000. Bases at chromosome 13 were often hypermethylated when dying; the mean methylation percentage difference was + 20.872%, compared to +0.125% across the whole genome (see also the increase for chromosome 13 in Fig. S12).

There were no genes that were differentially methylated in the CHG context.

For the CHG context there were 7167 hits for immune genes with differential methylation data in at least three samples, with a mean methylation difference of 0.019% (min = -1.645, max = 1.739). Again, the mean is near zero but significantly different due to the large number of genes (t_7302_ = 6.498, p < 0.001, 95% CIs = 0.013 to 0.025).

### CHH

Three colonies showed an increase in genome wide methylation rate, while two showed a decrease (Fig. 2c). Overall, the change was not different from zero (Wilcoxon signed-rank test, V = 6, p = 0.813). Chromosomes showed no clear change in median values from alive to dying (Wilcox-test, V = 1053, p = 0.269; Fig. S13). Variance of methylation rates at the chromosome level was similar between alive and dying colonies (variance alive = 3.092 x10^-10^, variance dying = 3.673 x10^-10^, F-test, F = 0.842, df = 69, p = 0.476). Groups of 1000 bases showed no clear change in median values from alive to dying (Wilcox-test, V = 1.073x10^14^, p = 0.477).Variance at the level of 1000 base pairs was also similar between alive and dying colonies (variance alive = 2.259x10^-6^, variance dying = 2.032x10^-6^, F-test, F = 1.112, df = 14,644457, p < 0.001; note the p value is very small due to the large sample size rather than a large effect size).

No cytosines were differentially methylated after correcting for multiple testing in the CHH context (Fig. 3e). There was however a clear spike in the % differences in methylation at the start of chromosome 13, in the same region as for the CHG context (Fig. 3f), but there was no equivalent spike in p-values (Fig. 3e). Cytosines on chromosome 13 were not consistently hypo- or hypermethylated; the mean methylation difference was -0.104% compared to +0.054% overall (which does not mark it out among the other chromosomes; see Fig. S13).

No gene-level analysis was possible for the CHH context (including for putative immune genes) as we could not calculate weighted methylation levels in this context for each gene, possibly due to the lack of overlap between cytosines methylated in the CHH context and cytosines appearing in gene bodies.

## Discussion

When comparing the genomes of social spiders dying from infection with the genomes of spiders from the same nests while healthy, we saw some increases in the mean and variance of genome- and chromosome-wide methylation levels, but these were only significant in the CHG context. At the per-base level, we saw few differentially methylated-bases in the CpG context, which were mostly associated with regions of repetitive DNA (although this was not an overrepresentation as more than half the genome consists of repeats), and none in the CHG and CHH contexts. There was however a spike of methylation changes in the CHG context in a particular region of chromosome 13. One gene was differentially methylated in the CpG context, but none were in the CHG context and we could not test this in the CHH context. Finally, genes with a known immune function in spiders had cytosines with mean methylation levels close to zero, suggesting this epigenetic modification is not key for changes in gene expression underpinning phenotypic immune responses.

Mean methylation levels in healthy colonies were around 12%, 0.009%, and 0.010% in the CpG, CHG, and CHH contexts respectively, and typically these each increased small amounts when the colonies were dying of infection. However, there was considerable among-nest variation in both means and changes, and so in no context were these differences statistically clear. We had predicted an increase in methylation levels, as we expected genes associated with an immune response to be upregulated and, in this species, methylation in the gene body is associated with higher expression (Liu *et al*., 2019). At the level of chromosomes and 1000 bases the increases in some of the contexts were clearer, suggesting the lack of clear increase at the nest level may have been due to the low number of colonies (five) rather than by a lack of effect. Similar results have been found in mosquitos *Aedes aegypti*, where infection with Wolbachia was associated with relatively few consistent changes in DNA methylation across the genome (Ye *et al*., 2013). In contrast, Abudukadier *et al*. (2021) found that Wolbachia infection in the spider *Hylyphantes graminicola* lead to genome-wide *reductions* in DNA methylation (due to reduced expression of the DNMT1 enzyme, see also Negri *et al*., 2009). Therefore, whether DNA methylation plays a key role in arthropod plastic immune response remains uncertain.

Research into the role of DNA methylation in phenotypic plasticity has typically focused on methylation in the CpG context, as it is more common than the other contexts. In mammals and plants methylation in the CpG context is clearly associated with changes in gene expression (He *et al*., 2011; Elhamamsy, 2016), but in invertebrates the association between DNA methylation and gene expression is much more variable (Richard *et al*., 2021; Duncan *et al*., 2022). Although not statistically significant after correcting for false discovery rate, patterns of differential methylation in the CHG context (and to a lesser extent in the CHH context) were much more localised than in the CpG context, suggesting this context may have a functional role in *S. dumicola*. Intriguingly, recent work in plants has suggested that differential methylation in non-CpG contexts may be associated with environmental variation and response to stress (Li *et al*., 2020; López *et al*., 2022; Antro *et al*., 2022). For example, woodland strawberry (*Fragaria vesca*) show global decrease of 1.8% in CHG methylation in response to salt stress and an increase of 3.1% when challenged with simulated hormone stress, while differential methylation was more common in the CHH context than either of the CpG or CHG contexts (López *et al*., 2022). While known to play important roles in plants (Kenchanmane Raju *et al*., 2019), the role of non-CpG context methylation in animals is not yet clear (although non-CpG methylation may play a role in the neural systems of birds and mammals; De Mendoza *et al*., 2021; Klughammer *et al*., 2023), but these recent findings along with our results highlight the need for more work to be done on these contexts.

We found mean methylation percentages changes of cytosines in putative immune genes to be around zero, in contrast to our predictions that changes in methylation would be targeted at these genes to alter their expression as part of the immune response. This supports our findings of limited base-level differential expression and overall suggests limited role of DNA methylation in controlling any plastic responses specifically to infection. In our gene-level analysis we identified one gene that was differentially methylated in the CpG context. Given the number of tests making conclusions about this gene would be unwise. Some studies on invertebrates have found that particular genes are differentially methylated in response to infection, such as genes associated with membrane transport in *A. aegypti* (Ye *et al*., 2013) and genes associated with the spliceosome, RNA transport, and protein processing in *B. mori* (Huang *et al*., 2019; see also: Kausar *et al*., 2022). However, given that methylation is not consistently associated with gene expression in insects (Richard *et al*., 2021; Duncan *et al*., 2022), these might be the exceptions rather than the rule.

Although across most of the genome we saw limited changes due to infection, there was a region on chromosome 13 that showed patterns of hypermethylation when infected in both the CHG and CHH contexts (although due to the number of bases tested none of these passed the threshold for statistical significance after correcting for false discovery rate). This was near the start of the chromosome, from approximately bases 704,000 to 821,000. Further analysis showed hypermethylation in this region was similar across genomic features (mean increases of 24.5% in introns and 20.5% in exons) which suggests a non-targeted increase rather directed methylation in order to alter gene expression. We use BLAST to search the NCBI database with the sequences of the two sequences showing the most clear (lowest p-value) change in methylation (p values < 3.00x10^-5^, compared to > 1.43x10^-2^ for all other genes; accession numbers in Table S3). The best match for one sequence was an uncharacterised protein also found in the bark scorpion (*Centruroides sculpturatus*), while the best match for the other was for a fructose aldolase from the bacterium *Mycoplasma anserisalpingitidis*. These hits do not suggest the region of hypermethylation in response to infection we have identified here plays much of a role in the immune system. Given there was also a spike in CHH methylation, an increase in untargeted methylation may have occurred at this region, although why that would be associated with the transition from healthy to infected state is unclear.

In conclusion, we found limited role for specific patterns of DNA methylation in the response to an immune challenge in social spiders *S. dumicola*. Instead, we saw some small genome-wide increases in the mean and variability of each type of methylation. These results suggest that there may be a genome-wide response in DNA methylation to infection, although whether this is adaptive plasticity or a neutral or maladaptive reaction to being infected is not clear. Further work on the prevalence and function of methylation in non-CpG contexts is required to better understand the intriguing patterns of differential methylation in the CHG context at a highly localised site in chromosome 13 we detected.

## Acknowledgements

We are grateful for the issued permissions to perform field work (permit number 1362/2017) granted by the Ministry of Environment and Tourism in Windhoek, Namibia. We thank those integral to the collection of the field data, especially Tharina Bird, Virginia Settepani, and Tom Tregenza. DF was supported by the Royal Society of Edinburgh (RSE Saltire Early Career Fellowship, grant no. 1940), while TB was supported by The Danish Council for Independent Research (grant no. 0135-00201B) and the Novo Nordisk Foundation (Interdisciplinary Synergy grant, grant no. NNF16OC0021110).

## References

Aagaard A, Liu S, Tregenza T, Braad Lund M, Schramm A, Verhoeven KJF, et al. (2022). Adapting to climate with limited genetic diversity: Nucleotide, DNA methylation and microbiome variation among populations of the social spider Stegodyphus dumicola. Molecular Ecology 31: 5765–5783.

Abudukadier A, Huang X, Peng Y, Zhang F, Liu H, Chen J, et al. (2021). The potential association between Wolbachia infection and DNA methylation in Hylyphantes graminicola (Araneae: Linyphiidae). Symbiosis 83: 183–191.

Akalin A, Kormaksson M, Li S, Garrett-Bakelman FE, Figueroa ME, Melnick A, et al. (2012). MethylKit: a comprehensive R package for the analysis of genome-wide DNA methylation profiles. Genome Biology 13: 87.

Altermatt F, Ebert D (2008). Genetic diversity of Daphnia magna populations enhances resistance to parasites. Ecology Letters 11: 918–928.

Antro MV, Prelovsek S, Ivanovic S, Gawehns F, Wagemaker NCAM, Mysara M, et al. (2022). DNA methylation in clonal Duckweed lineages (Lemna minor L.) reflects current and historical environmental exposures.: 2022.08.23.504803.

Bechsgaard J, Jorgensen TH, Jønsson AK, Schou M, Bilde T (2022). Impaired immune function accompanies social evolution in spiders. Biology Letters 18: 20220331.

Bechsgaard J, Vanthournout B, Funch P, Vestbo S, Gibbs RA, Richards S, et al. (2016). Comparative genomic study of arachnid immune systems indicates loss of beta-1,3-glucanase-related proteins and the immune deficiency pathway. Journal of Evolutionary Biology 29: 277–291.

Bird A (2002). DNA methylation patterns and epigenetic memory. Genes & Development 16: 6–21.

Busck MM, Lund MB, Bird TL, Bechsgaard JS, Bilde T, Schramm A (2022). Temporal and spatial microbiome dynamics across natural populations of the social spider Stegodyphus dumicola. FEMS Microbiology Ecology 98.

Busck MM, Settepani V, Bechsgaard J, Lund MB, Bilde T, Schramm A (2020). Microbiomes and Specific Symbionts of Social Spiders: Compositional Patterns in Host Species, Populations, and Nests. Frontiers in Microbiology 11: 1845.

Cardoso-Júnior CAM, Yagound B, Ronai I, Remnant EJ, Hartfelder K, Oldroyd BP (2021). DNA methylation is not a driver of gene expression reprogramming in young honey bee workers. Molecular Ecology 30: 4804–4818.

Chen Z, Boehnke M, Wen X, Mukherjee B (2021). Revisiting the genome-wide significance threshold for common variant GWAS. G3 Genes|Genomes|Genetics 11.

Claudio-Piedras F, Recio-Tótoro B, Condé R, Hernández-Tablas JM, Hurtado-Sil G, Lanz-Mendoza H (2020). DNA Methylation in Anopheles albimanus Modulates the Midgut Immune Response Against Plasmodium berghei. Frontiers in Immunology 10: 3025.

Cokus SJ, Feng S, Zhang X, Chen Z, Merriman B, Haudenschild CD, et al. (2008). Shotgun bisulphite sequencing of the Arabidopsis genome reveals DNA methylation patterning. Nature 2008 452:7184 452: 215–219.

Cremer S, Armitage SAO, Schmid-Hempel P (2007). Social Immunity. Current Biology 17: R693–R702.

De Mendoza A, Poppe D, Buckberry S, Pflueger J, Albertin CB, Daish T, et al. (2021). The emergence of the brain non-CpG methylation system in vertebrates. Nat Ecol Evol 5: 369–378.

Deleris A, Halter T, Navarro L (2016). DNA Methylation and Demethylation in Plant Immunity. Annual Review of Phytopathology 54: 579–603.

Dubin MJ, Zhang P, Meng D, Remigereau MS, Osborne EJ, Casale FP, et al. (2015). DNA methylation in Arabidopsis has a genetic basis and shows evidence of local adaptation. eLife 4.

Duncan EJ, Cunningham CB, Dearden PK (2022). Phenotypic Plasticity: What Has DNA Methylation Got to Do with It? Insects 13.

Elhamamsy AR (2016). DNA methylation dynamics in plants and mammals: overview of regulation and dysregulation. Cell Biochemistry and Function 34: 289–298.

Flores KB, Wolschin F, Amdam GV (2013). The role of methylation of DNA in environmental adaptation. Integrative and Comparative Biology 53: 359–372.

Hämälä T, Ning W, Kuittinen H, Aryamanesh N, Savolainen O (2022). Environmental response in gene expression and DNA methylation reveals factors influencing the adaptive potential of Arabidopsis lyrata. bioRxiv: 2022.03.24.485516.

Hansen KD, Langmead B, Irizarry RA (2012). BSmooth: from whole genome bisulfite sequencing reads to differentially methylated regions. Genome Biology 13: 1–10.

Hatchwell BJ (2010). Cryptic kin selection: Kin structure in vertebrate populations and opportunities for kin-directed cooperation. Ethology 116: 203–216.

He XJ, Chen T, Zhu JK (2011). Regulation and function of DNA methylation in plants and animals. Cell Research 2011 21:3 21: 442–465.

Huang H, Wu P, Zhang S, Shang Q, Yin H, Hou Q, et al. (2019). DNA methylomes and transcriptomes analysis reveal implication of host DNA methylation machinery in BmNPV proliferation in Bombyx mori. BMC Genomics 20: 736.

Kausar S, Liu R, Gul I, Abbas MN, Cui H (2022). Transcriptome Sequencing Highlights the Regulatory Role of DNA Methylation in Immune-Related Genes’ Expression of Chinese Oak Silkworm, Antheraea pernyi. Insects 13: 296.

Kenchanmane Raju SK, Ritter EJ, Niederhuth CE (2019). Establishment, maintenance, and biological roles of non-CG methylation in plants. Essays in Biochemistry 63: 743–755.

Klughammer J, Romanovskaia D, Nemc A, Posautz A, Seid CA, Schuster LC, et al. (2023). Comparative analysis of genome-scale, base-resolution DNA methylation profiles across 580 animal species. Nat Commun 14: 232.

Krueger F, Andrews SR (2011). Bismark: a flexible aligner and methylation caller for Bisulfite-Seq applications. Bioinformatics 27: 1571–1572.

Kuhn-Nentwig L, Nentwig W (2013). The immune system of spiders. In: Spider Ecophysiology, Springer-Verlag Berlin Heidelberg, pp 81–91.

Lawrence M, Huber W, Pagès H, Aboyoun P, Carlson M, Gentleman R, et al. (2013). Software for Computing and Annotating Genomic Ranges. PLOS Computational Biology 9: e1003118.

Ledon-Rettig CC, Richards CL, Martin LB (2012). Epigenetics for behavioral ecologists. Behavioral Ecology 24: 311–324.

Lee S, Cook D, Lawrence M (2019). Plyranges: A grammar of genomic data transformation. Genome Biology 20: 1–10.

Li R, Hu F, Li B, Zhang Y, Chen M, Fan T, et al. (2020). Whole genome bisulfite sequencing methylome analysis of mulberry (Morus alba) reveals epigenome modifications in response to drought stress. Sci Rep 10: 8013.

Libbrecht R, Oxley PR, Keller L, Kronauer DJC (2016). Robust DNA methylation in the clonal raider ant brain. Current Biology 26: 391–395.

Liu S, Aagaard A, Bechsgaard J, Bilde T (2019). DNA methylation patterns in the social spider, Stegodyphus dumicola. Genes 10: 137.

López M-E, Roquis D, Becker C, Denoyes B, Bucher E (2022). DNA methylation dynamics during stress response in woodland strawberry (Fragaria vesca). Horticulture Research 9: uhac174.

Lubin Y, Bilde T (2007). The Evolution of Sociality in Spiders. Advances in the Study of Behavior 37: 83–145.

Ma J, Bechsgaard J, Aagaard A, Villesen P, Bilde T, Schierup M (2024). Sociality in spiders is an evolutionary dead-end.

Majer M, Svenning JC, Bilde T (2013). Habitat productivity constrains the distribution of social spiders across continents - case study of the genus Stegodyphus. Frontiers in Zoology 10: 1–10.

Marshall H, Lonsdale ZN, Mallon EB (2019). Methylation and gene expression differences between reproductive and sterile bumblebee workers. Evolution Letters 3: 485–499.

McCullagh P, Nelder JA (1989). Generalized Linear Models. Chapman & Hall/CRC: New York.

McEwen BL, Lichtenstein JLL, Fisher DN, Wright CM, Chism GT, Pinter-Wollman N, et al. (2020). Predictors of colony extinction vary by habitat type in social spiders. Behavioral Ecology and Sociobiology 74.

Morales-Nebreda L, McLafferty FS, Singer BD (2019). DNA methylation as a transcriptional regulator of the immune system. Translational Research 204: 1–18.

Mostoslavsky R, Bergman Y (1997). DNA methylation: regulation of gene expression and role in the immune system. Biochimica et Biophysica Acta 1333: 1997–2026.

Negri I, Franchini A, Gonella E, Daffonchio D, Mazzoglio PJ, Mandrioli M, et al. (2009). Unravelling the Wolbachia evolutionary role: the reprogramming of the host genomic imprinting. Proc R Soc B 276: 2485–2491.

Nunn CL, Jordán F, McCabe CM, Verdolin JL, Fewell JH (2015). Infectious disease and group size: more than just a numbers game. Philosophical transactions of the Royal Society of London Series B, Biological sciences 370: 20140111-.

Palmer WJ, Jiggins FM (2015). Comparative Genomics Reveals the Origins and Diversity of Arthropod Immune Systems. Molecular Biology and Evolution 32: 2111–2129.

Richard G, Jaquiéry J, Le Trionnaire G (2021). Contribution of Epigenetic Mechanisms in the Regulation of Environmentally-Induced Polyphenism in Insects. Insects 2021, Vol 12, Page 649 12: 649.

Risch N, Merikangas K (1996). The future of genetic studies of complex human diseases. Science 273: 1516–1517.

Rose C, Lund MB, Søgård AM, Busck MM, Bechsgaard JS, Schramm A, et al. (2023). Social transmission of bacterial symbionts homogenizes the microbiome within and across generations of group-living spiders. ISME Communications 3: 60.

Schmid-Hempel P (1988). Parasites in Social Insects (P Schmid-Hempel, Ed.). Princeton University Press: Princeton.

Schultz MD, Schmitz RJ, Ecker JR (2012). ‘Leveling’ the playing field for analyses of single-base resolution DNA methylomes. Trends in Genetics 28: 583–585.

Settepani V, Schou MF, Greve M, Grinsted L, Bechsgaard J, Bilde T (2017). Evolution of sociality in spiders leads to depleted genomic diversity at both population and species levels. Molecular Ecology 26: 4197–4210.

Tong C, Najm GM, Pinter-Wollman N, Pruitt JN, Linksvayer TA, Pisani D (2020). Comparative Genomics Identifies Putative Signatures of Sociality in Spiders. Genome Biology and Evolution 12: 122–133.

Van Valen L (1973). A new evolutionary law. Evolutionary Theory 1: 1–30.

Wu P, Jie W, Shang Q, Annan E, Jiang X, Hou C, et al. (2017). DNA methylation in silkworm genome may provide insights into epigenetic regulation of response to Bombyx mori cypovirus infection. Scientific Reports 7.

Ye YH, Woolfit M, Huttley GA, Rancès E, Caragata EP, Popovici J, et al. (2013). Infection with a Virulent Strain of Wolbachia Disrupts Genome Wide-Patterns of Cytosine Methylation in the Mosquito Aedes aegypti (K Bourtzis, Ed.). PLoS ONE 8: e66482.

