## Supplementary materials for "Exploring changes in social spider DNA methylation profiles when succumbing to infection in CpG, CHG, and CHH contexts"

### Tables

**Table S1**. Sample details for each nest used in the study, including overall cytosine information before and after duplication, and split by context (CpG, CHG, and CHH). Two or three siders from the same nest sampled on the same day were pooled for whole-genome bisulfite sequencing, giving ten samples in total. See Busck *et al*. (2022) for further details on sample collection and other information. Note that values given here are directly from the Bismarck extraction summary, not from the binomial test we used to assign methylation status to a cytosine base, and so the percentages given may not match those in the text exactly.

| Sampling date | Nest ID | Status | Before de-duplication | | After de-duplication | | | | |
| --- | --- | --- | --- | --- | --- | --- | --- | --- | --- |
|  |  |  | Methylation call strings | Total cytosines | Methylation call strings | Number of cytosines | Total cytosines methylated | Number cytosines unmethylated | % methylated |
| 06/07/2017 | T052 | Alive | 499939956 | 6998875167 | 489460270 | 6852752125 | 160031413 | 6692720712 | 2.34 |
| 01/01/2018 | T052 | Dying | 487489320 | 6744275209 | 477767950 | 6611086366 | 160650910 | 6450435456 | 2.43 |
| 06/07/2017 | T053 | Alive | 497255288 | 6957647569 | 487549728 | 6822425494 | 158966537 | 6663458957 | 2.33 |
| 30/06/2018 | T053 | Dying | 484499766 | 6768031872 | 474304178 | 6627009068 | 159381842 | 6467627226 | 2.41 |
| 06/07/2017 | T054 | Alive | 517470788 | 7302649813 | 505546100 | 7134626447 | 164273212 | 6970353235 | 2.30 |
| 01/01/2018 | T054 | Dying | 503233506 | 7296325886 | 491490396 | 7126064315 | 160780667 | 6965283648 | 2.26 |
| 13/04/2017 | T055 | Alive | 507019726 | 6906365487 | 496613140 | 6767133436 | 153552317 | 6613581119 | 2.27 |
| 11/10/2017 | T055 | Dying | 514884040 | 7340143857 | 503994770 | 7185047245 | 170508707 | 7014538538 | 2.37 |
| 13/04/2017 | T056 | Alive | 485743712 | 6749701621 | 474434188 | 6592062927 | 146189027 | 6445873900 | 2.22 |
| 11/10/2017 | T056 | Dying | 488214012 | 7026369819 | 472768156 | 6800860305 | 151492421 | 6649367884 | 2.23 |

**Table S1**. continued

| Sampling date | Nest ID | Status | CpG context | | | | CHG context | | | | CHH context | | | |
| --- | --- | --- | --- | --- | --- | --- | --- | --- | --- | --- | --- | --- | --- | --- |
|  |  |  | Total Cs | Methylated Cs | Unmethylated Cs | % methylated | Total Cs | Methylated Cs | Unmethylated Cs | % methylated | Total Cs | Methylated Cs | Unmethylated Cs | % methylated |
| 06/07/2017 | T052 | Alive | 1125393564 | 117387342 | 1008006222 | 10.43 | 1248282543 | 8928833 | 1239353710 | 0.72 | 4479076018 | 33715238 | 4445360780 | 0.75 |
| 01/01/2018 | T052 | Dying | 1101990486 | 110862966 | 991127520 | 10.06 | 1207262980 | 10150682 | 1197112298 | 0.84 | 4301832900 | 39637262 | 4262195638 | 0.92 |
| 06/07/2017 | T053 | Alive | 1124868318 | 114539021 | 1010329297 | 10.18 | 1246870703 | 9059417 | 1237811286 | 0.73 | 4450686473 | 35368099 | 4415318374 | 0.79 |
| 30/06/2018 | T053 | Dying | 1106356858 | 115386566 | 990970292 | 10.43 | 1211477079 | 9262935 | 1202214144 | 0.76 | 4309175131 | 34732341 | 4274442790 | 0.81 |
| 06/07/2017 | T054 | Alive | 1161850858 | 118779149 | 1043071709 | 10.22 | 1292831413 | 9395758 | 1283435655 | 0.73 | 4679944176 | 36098305 | 4643845871 | 0.77 |
| 01/01/2018 | T054 | Dying | 1155749146 | 121080562 | 1034668584 | 10.48 | 1279689597 | 8222858 | 1271466739 | 0.64 | 4690625572 | 31477247 | 4659148325 | 0.67 |
| 13/04/2017 | T055 | Alive | 1113166479 | 109962759 | 1003203720 | 9.88 | 1232547014 | 9038403 | 1223508611 | 0.73 | 4421419943 | 34551155 | 4386868788 | 0.78 |
| 11/10/2017 | T055 | Dying | 1169489260 | 120916823 | 1048572437 | 10.34 | 1304746164 | 9947179 | 1294798985 | 0.76 | 4710811821 | 39644705 | 4671167116 | 0.84 |
| 13/04/2017 | T056 | Alive | 1076161166 | 106597098 | 969564068 | 9.91 | 1193379595 | 7989516 | 1185390079 | 0.67 | 4322522166 | 31602413 | 4290919753 | 0.73 |
| 11/10/2017 | T056 | Dying | 1109505417 | 113606863 | 995898554 | 10.24 | 1222916152 | 7911310 | 1215004842 | 0.65 | 4468438736 | 29974248 | 4438464488 | 0.67 |

**Table S2**. FlyBase ID numbers for the 46 genes with putative immune function which we used for our BLAST

FBgn0000229, FBgn0000250, FBgn0003326, FBgn0003495, FBgn0004429, FBgn0004431, FBgn0010303, FBgn0010441, FBgn0014018, FBgn0016917, FBgn0020381, FBgn0024222, FBgn0026323, FBgn0027594, FBgn0029114, FBgn0030164, FBgn0031959, FBgn0032362, FBgn0033159, FBgn0033327, FBgn0033367, FBgn0033402, FBgn0035056, FBgn0035379, FBgn0035813, FBgn0036494, FBgn0037906, FBgn0038928, FBgn0039016, FBgn0041180, FBgn0041181, FBgn0041182, FBgn0041205, FBgn0043575, FBgn0043578, FBgn0043903, FBgn0050281, FBgn0087011, FBgn0260632, FBgn0261363, FBgn0262473, FBgn0262739, FBgn0262870, FBgn0263219, FBgn0267488, FBgn0283531

**Table S3**. Accession numbers of the best match in the NCBI database for the coding sequences of the two genes in the region of chromosome 13 showing hypermethylation in the CHG context.

| XP_023212243.1 | WP_146367770.1 |
| --- | --- |

### Figures

[see attached zip folder]

**Figures S1-10**. Plots of the mean percentage methylation in CpG, CHG, and CHH contexts for reads 1 and 2 across the 100 base pairs of each segment. These are known as “M-bias” plots and are used to determine how much of each end of each read to discard before aligning to the genome. We see high variability in the first five bases of the 5’ prime end (bases 1-5 in read 1) and in the first two bases of the 3’ end (bases 1-2 in read 2) and so we removed these. Plots feature the five colonies (T52, T53, T54, T55, and T56) at each of the “alive” and “dying” time points.

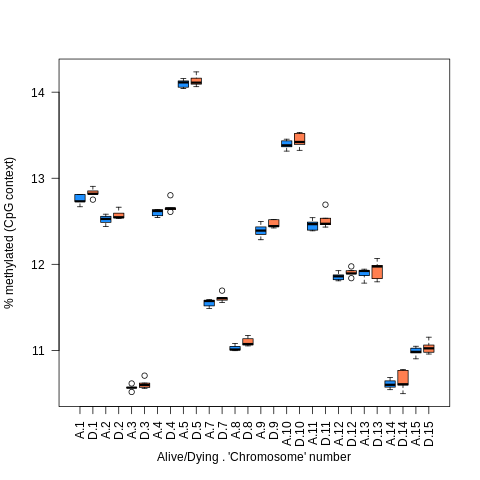

**Figure S11**. Plot of percentage of cytosines methylated in CpG context per chromosome, between alive (blue) and dying (orange) time points. Each box and whisker represents the five data points from each nest. Note “chromosomes” 6 and 16 are not shown as they are scaffolds that are not true chromosomes.

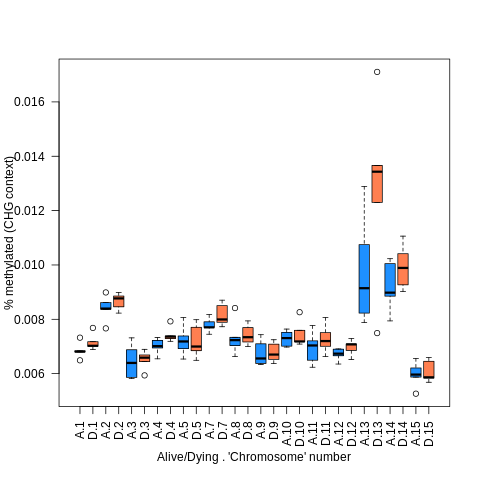

**Figure S12**. Plot of percentage of cytosines methylated in CHG context per chromosome, between alive (blue) and dying (orange) time points. Each box and whisker represents the five data points from each nest. Note “chromosomes” 6 and 16 are not shown as they are scaffolds that are not true chromosomes.

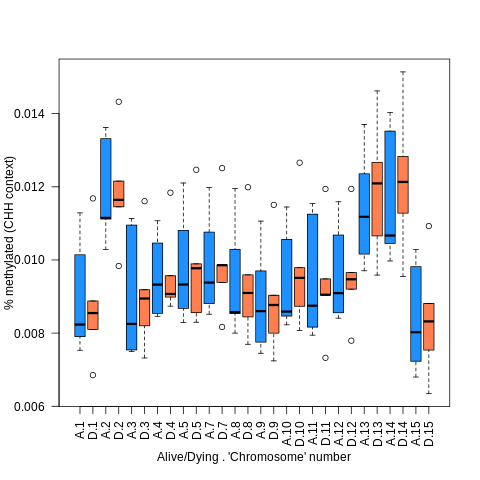

**Figure S13**. Plot of percentage of cytosines methylated in CHH context per chromosome, between alive (blue) and dying (orange) time points. Each box and whisker represents the five data points from each nest. Note “chromosomes” 6 and 16 are not shown as they are scaffolds that are not true chromosomes.
